# Curation of the Deep Green list of unannotated green lineage proteins to enable structural and functional characterization

**DOI:** 10.1101/2022.09.30.510186

**Authors:** Eric P. Knoshaug, Peipei Sun, Ambarish Nag, Huong Nguyen, Erin M. Mattoon, Ningning Zhang, Jian Liu, Chen Chen, Jianlin Cheng, Ru Zhang, Peter St. John, James Umen

**Affiliations:** Biosciences Center, National Renewable Energy Laboratory, Golden CO 80401; Donald Danforth Plant Science Center, St. Louis MO 63132, USA; Computational Sciences Center, National Renewable Energy Laboratory, Golden CO 80401; Plant and Microbial Biosciences Program, Division of Biology and Biomedical Sciences, Washington University in Saint Louis, St. Louis, MO 63130, USA; Institute of Genomics for Crop Abiotic Stress Tolerance, Department of Plant and Soil Science, Texas Tech University, Lubbock, TX 79409, USA; Department of Electrical Engineering and Computer Science, University of Missouri, Columbia, MO 65211, USA

**Keywords:** protein structure, functional annotation, Deep Green conserved proteins, Arabidopsis, Setaria

## Abstract

An explosion of sequenced genomes and predicted proteomes enabled by low cost deep sequencing has revolutionized biology. Unfortunately, protein functional annotation is more complex, and has not kept pace with the sequencing revolution. We identified unannotated proteins in three model organisms representing distinct parts of the green lineage (Viridiplantae); *Arabidopsis thaliana* (dicot), *Setaria viridis* (monocot), and *Chlamydomonas reinhardtii* (Chlorophyte alga). Using similarity searching we found the subset of unannotated proteins that were conserved between these species and defined them as Deep Green proteins. Informatic, genomic, and structural predictions were leveraged to begin inferring functional information about Deep Green genes and proteins. The Deep Green set was enriched for proteins with predicted chloroplast targeting signals that are predictive of photosynthetic or plastid functions. Strikingly, structural predictions using AlphaFold and comparisons to known structures show that a significant proportion of Deep Green proteins may possess novel protein tertiary structures. The Deep Green genes and proteins provide a starting resource of high value targets for further investigation of potentially new protein structures and functions that are conserved in the green lineage.

## Introduction

The genome sequencing revolution of the past two decades has removed a major barrier to identifying and describing the genetic toolkits used by green lineage organisms for growth, photosynthesis, and adaptation to diverse conditions. The number of sequenced plant genomes is growing exponentially, but the resources for comprehensive, experimental structural and functional protein annotation are lagging behind (Ellens et al., 2017). Homology-based annotations are a simple and rapid means of predicting protein function in the absence of any other functional data. In homology-based annotations, new sequences are searched for similarity to proteins in other species, some of which may already have known functions that may be assigned to the newly predicted protein “by proxy”. It is generally assumed that conservation at the sequence level implies conservation of function, but this correlation is not perfect (Blaby-Hass et al., 2011). New sequences can also be searched for the presence of conserved domains using sensitive Hidden Markov Models (HMM) and/or multiple sequence alignments (Soding, 2005). The Protein ANalysis THrough Evolutionary Relationships (PANTHER), Protein Families (Pfam), and Clusters of Orthologous Groups of proteins (COG) and the eukaryote-specific version EuKaryotic Orthologous Groups (KOG) are powerful classification tools that leverage growing sets of data to improve annotation based on sequence similarity (Tatusov et al., 2003, Koonin et al., 2004, Mi et al., 2012, Finn et al., 2016, Mi et al., 2016, Bolger et al., 2017, El-Gebali et al., 2018). The above approaches will identify functions and/or structural domains for around half of all proteins in newly sequenced plant genomes, with some variation dependent on genome size and complexity as well as taxonomic position (Hanson et al., 2010). Even well annotated genomes from budding yeast and humans still contain approximately 30% unannotated proteins, and 30-40% of these unannotated proteins (~10% total) are likely to have an uncharacterized catalytic function [1]. Likewise, every genome contains predicted proteins that are unannotated because they either have no similarity to characterized proteins or have limited information available beyond the possible presence or absence of domains. Unannotated proteins have typically accounted for approximately 40-60% in plants and algae genomes (Berardini et al., 2015, Niehaus et al., 2015, Blaby-Haas et al., 2019) with functional assignments slowly increasing.

The potential for discovery of new structures, catalytic activities, and biological functions among unknown proteins is enormous, but these proteins also represent a huge challenge due to their overwhelming numbers which increase whenever a new genome is sequenced (Fox et al., 2008, Hanson et al., 2010). Having improved functional information for plant proteins is especially important as the need for faster deployment of biotechnology and breeding resources is essential to continue the pace of development of agriculture and bioenergy crops to maintain food and energy security in a challenging global environment with an increasing human population and growing climate instability. An integral part of overcoming challenge will involve crops with improved tolerance for abiotic stresses such as temperature, light, salt, and nutrient deficiency (Ahanger et al., 2017, Maggio et al., 2018, Varshney et al., 2018).

Functional annotations of plant proteins lags behind that for animals, fungi, and prokaryotes due to fewer resources devoted to plant research and incomplete plant genome annotation, though this situation is improving with dedicated portals and tool development on platforms such as Phytozome, PLAZA, and Gramene (Goodstein et al., 2012, Proost et al., 2015, Tello-Ruiz et al., 2018). Currently, approximately 12% of predicted proteins have an experimentally determined structure or structural data derived from related proteins in the Protein Data Bank (PDB) for *Arabidopsis thaliana*, compared to more than 25% in *Saccharomyces cerevisiae* and more than 30% in *Homo sapiens* (Callaway, 2022). One approach for prioritizing unknown/unannotated genes and proteins for further functional analyses is to demand sequence conservation across one or more taxa (aka phylogenomics). While this does not directly help with annotation, it ensures that information obtained about that gene or associated protein from one species will be impactful as it can be leveraged across species. Indeed, phylogenomics approaches have been used successfully to obtain new biological information in multiple contexts. One particularly useful deployment of phylogenomics defined GreenCut proteins, found in multiple photosynthetic taxa, but not in non-photosynthetic eukaryotes. The GreenCut set of unknown/uncharacterized proteins was first defined when annotating the genome of the green alga *Chlamydomonas reinhardtii* (Merchant et al., 2007), and has since been expanded and updated (Karpowicz et al., 2011). Validating its usefulness, many of the original unknowns in the GreenCut list were found to have key functions in photosynthetic processes (Arthur et al., 2019, Wakao et al., 2021). A limitation of the GreenCut list is that it spans a wide taxonomic breadth including non-green-lineage species such as diatoms, and red and brown algae and demands high similarity scores between candidates. This stringency ensures higher confidence of an important photosynthetic function for candidates but may exclude proteins that diverged too rapidly to detect similarities across large phylogenetic distances.

Here we took a comprehensive approach using inclusive criteria to create the Deep Green list of unannotated green lineage proteins. This list is based on identification and curation of conserved unknown proteins in three green lineages (i.e. Viridiplantae) species: *Chlamydomonas reinhardtii, Arabidopsis thaliana*, and *Setaria viridis*. Preliminary characterization of Deep Green proteins using in silico methods and published data sets is reported here and has revealed similarities among these proteins including enrichment in predicted chloroplast, photosynthetic, and stress related functions, and identified multiple predicted families of novel protein structures.

## Results and Discussion

Our objective was to define and begin characterizing a set of poorly annotated or unannotated conserved green lineage proteins. To do so we chose three focal species: *Arabidopsis thaliana* (Arabidopsis), *Setaria viridis* (Setaria), and *Chlamydomonas reinhardtii* (Chlamydomonas). Arabidopsis has the best studied and well annotated genome of any angiosperm species (Berardini et al., 2015), while Setaria (green foxtail millet) is an emerging model C4 grass and bioenergy feedstock model that has a small stature, short generation time, and is genetically tractable (Huang et al., 2016, Zhu et al., 2017, Hu et al., 2018). Together, Arabidopsis and Setaria represent two major branches of angiosperms, dicots and monocots, respectively. The unicellular green alga Chlamydomonas is a well-established model for investigating cellular processes in photosynthetic eukaryotes due to its fast growth rates, haploid genome, low levels of gene duplication, and availability of high-throughput genetics and genomics tools (Sasso et al., 2018).

### Identification of conserved protein families

To reduce search complexity and to help identify unannotated proteins, we developed a down-selection strategy (**Figure 1A**). The first step involved grouping the predicted proteins in each species into families with highly similar sequences (>30% sequence identity, >50% overlap), while at the same time merging annotations among family members (**Tables S1-S3**). This paralog grouping reduced sequence search space from 27654, 38334, and 17741 predicted primary proteins to 13058, 19951, and 14076 in Arabidopsis, Setaria, and Chlamydomonas respectively (**Figure 1B**), and ensured that unannotated members of a paralog family were not included when one of their family members was already annotated. The h3-cd-hit algorithm which was used for grouping into families also selected a lead protein in each cluster which served as a representative for similarity searching between proteomes of the three focal species (**Tables S4-S6**). For each protein family, Phytozome annotations were collected, merged, and used to identify unannotated or poorly annotated families in subsequent steps. Because our goal was to identify conserved unannotated proteins, the next step in the process was to find protein families that were shared between at least two of the three species. Using the basic local alignment search tool for protein (BLASTP) among lead proteins from each species, the top high-scoring segment pair (hsp) was identified and those which passed a similarity threshold (e-value cutoff threshold of 10^-3^; >40% of the sequence length aligned for at least one of the two proteins) were used to further group proteins into superfamilies with shared annotations (**Figure 2, Table S7**).

**Figure 1.**
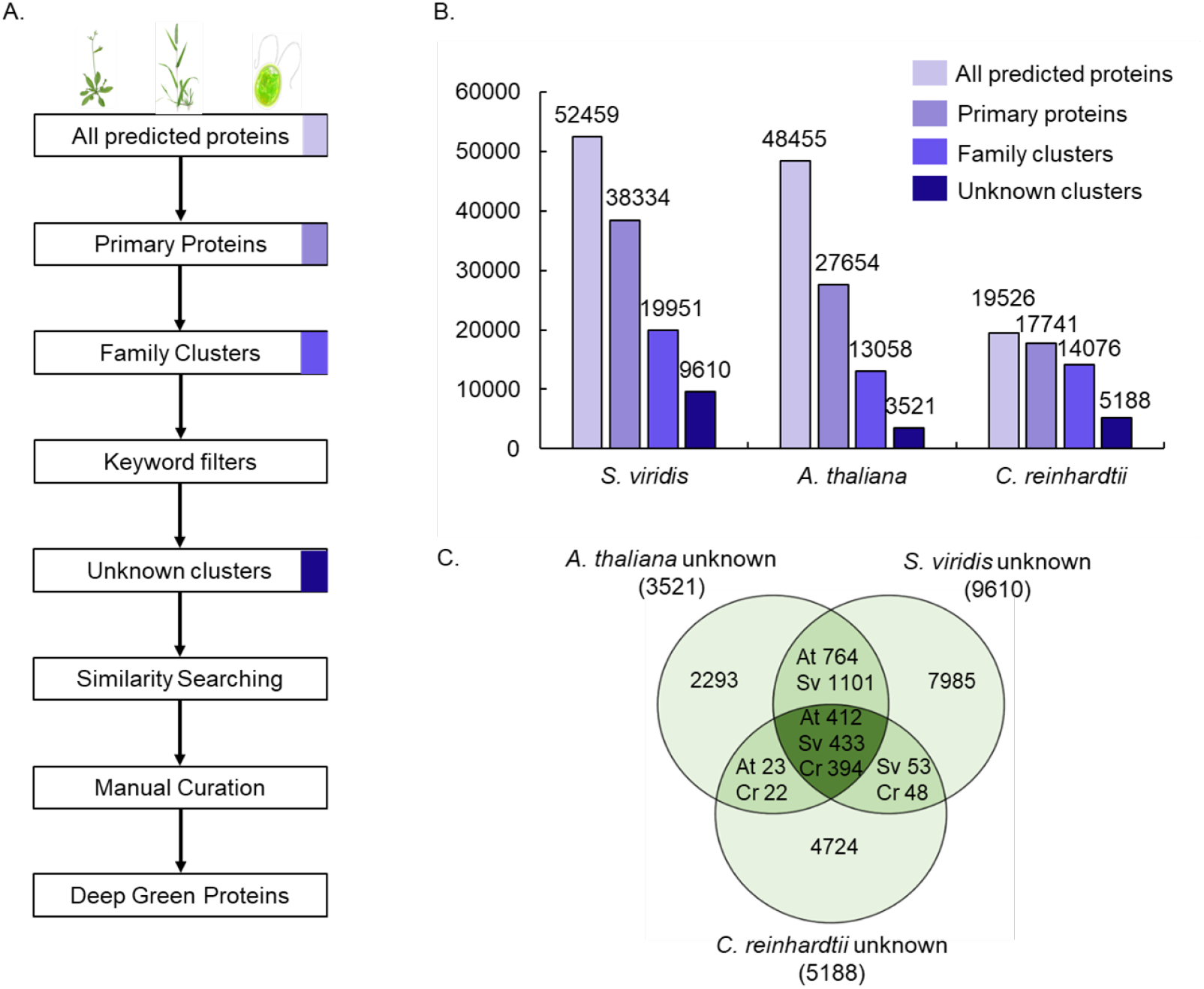
(A) Deep Green down-selection flowchart, (B) Number of proteins in each species at each of the down-selection steps, (C) Venn diagram showing overlaps between conserved proteins which define the Deep Green set.

**Figure 2.**
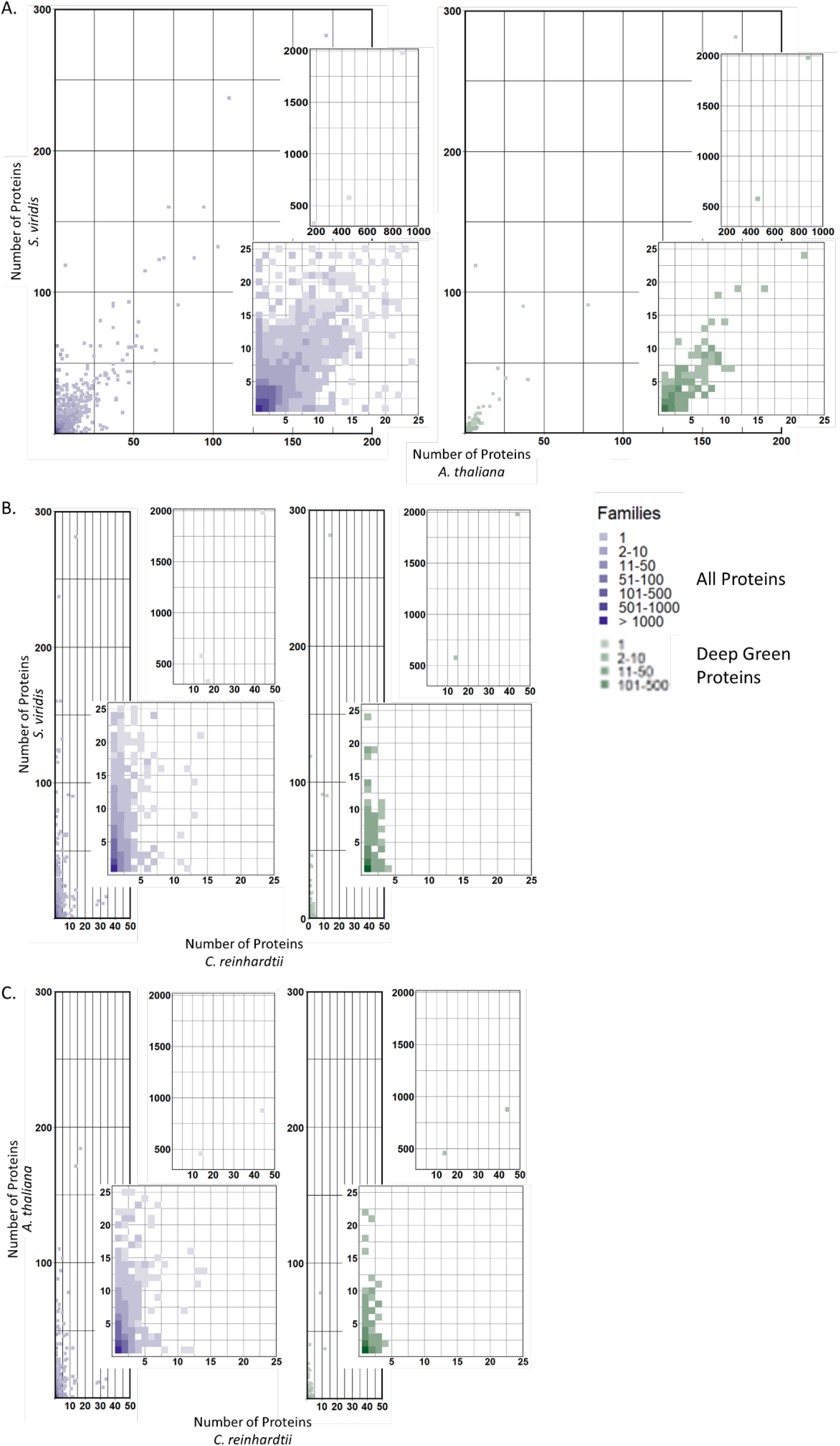
Comparison of protein family sizes in (A) Arabidopsis and Setaria, (B) Chlamydomonas and Setaria, and (C) Chlamydomonas and Arabidopsis. Insets show in greater detail families containing 25 or fewer proteins or those containing greater than 300 members. Axes indicate the number of proteins in a family and color scale denotes the number of families at each position in the graph. For example, the darkest point in each graph (1:1) represents >1000 protein families with a single family member in each of the two species.

### Identification of unannotated/unknown proteins

The families (**Tables S4-S6**) were filtered to select poorly annotated or unannotated members based on keyword searching of aggregated annotations (**Figure 1A**). If a family in one species was well annotated or characterized, its conserved counterparts in the other two species were also considered annotated and removed from consideration. If the definition line (defline) annotation was blank, or contained the terms “unknown”, “undefined”, “uncharacterized”, “hypothetical”, “domain unknown function”, “expressed protein”, “transmembrane”, “function unknown”, “predicted protein” and “conserved in plant or green lineage”, “anykrin”, “fbox”, “tetra- or penta-tricopeptide”, the family was retained. In Arabidopsis, a list of uncharacterized/unannotated proteins was recently released and incorporated into our unknown protein lists (phoenixbioinformatics.org, https://conf.phoenixbioinformatics.org/pages/viewpage.action?pageId=22807120.) The conserved unknown proteins within each list were further curated by manual inspection of annotations and by searching for the protein or gene IDs in publications. If a protein had been functionally characterized (e.g., published mutant phenotype) it was removed from the list. Manual searching for ambiguous or poor quality annotations among the lists of annotated clusters (e. g., Arabidopsis clusters 7885 and 9084) was also done to enable inclusion of proteins that did not fit the defline criteria above. In parallel we updated the GreenCut2 protein list with new gene IDs based on the Phytozome 5.5 genome assembly and by removing those in the list that had been characterized since the original GreenCut2 publication (Karpowicz et al., 2011, Arthur et al., 2019) and the remaining GreenCut2 proteins were merged into the final list of unannotated proteins. This final manual curation led to 3521, 9610, and 5188 uncharacterized or poorly characterized families in Arabidopsis, Setaria, and Chlamydomonas respectively (**Figure 1B**). Finally, we sorted the unknown protein lists to identify overlaps between each of the species (**Figure 1C, Tables S4-S6**) to define the final Deep Green protein list.

### Characterization of Unknown and Deep Green proteins

We interrogated the Deep Green list in several ways to provide preliminary information on potential functions. Deep Green proteins in all the three species were strongly enriched for chloroplast targeting and the plant members of this set were also de-enriched for nuclear targeting (**Figure 3**).

**Figure 3.**
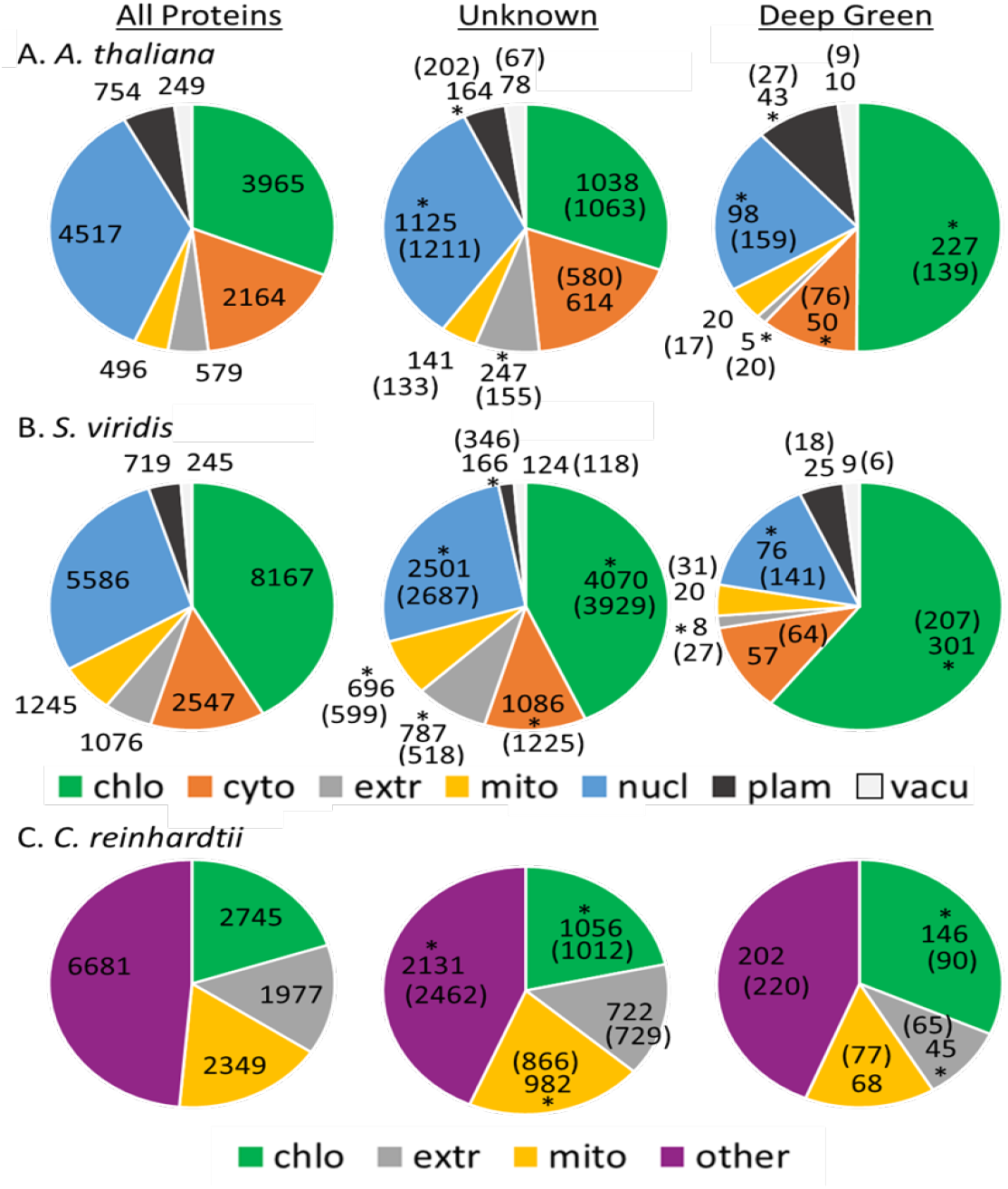
Predicted protein localization for (A) Arabidopsis, (B) Setaria, and (C) Chlamydomonas. Predictions for Arabidopsis and Setaria were done using WoLFPSORT. Predictions for Chlamydomonas were done using Predalgo. Numbers in parentheses for the Unknown and Deep Green protein sets are the expected number using Fisher’s Exact Test (background size 13058, 19952, and 14076 for the three species respectively) based on the total number of proteins and their predicted localization. Significant enrichment or de-enrichment (*FDR corrected p-value < 0.05) is indicated. Cellular localization abbreviations: chlo, chloroplast; cyto, cytosol; extr, extracellular; mito, mitochondria; nucl, nucleus; plam, plasma membrane; vacu, vacuole.

Importantly, the strong chloroplast localization enrichment for predicted proteins in each species was not seen in the set of all unknown proteins for each species where there was just a slight enrichment. These findings suggest that there is a significant number of conserved proteins with chloroplast functions that have yet to be characterized. Deep Green and unknown proteins were also significantly enriched in proteins predicted to contain one or more transmembrane domain(s) as compared to all predicted protein families (**Figure S1**).

We next performed co-expression analysis for Chlamydomonas Deep Green genes using published transcriptome data sets. An important resource was a previously described high-resolution diurnal data set for synchronized Chlamydomonas cultures (Zones et al., 2015). In that study, around 80% of genes with detectable expression (~12,000) showed strong periodic diurnal or cell-cycle-controlled expression patterns (**Figure 4A**). Genes coding for the Deep Green proteins showed higher overall average expression levels compared with all expressed genes of 3.63 versus 1.91 log2RPKM respectively (**Figure 4B**). We also investigated the distribution and enrichment of Deep Green genes in 18 diurnal clustered and unclustered expression groups and found them to be significantly over-represented in clusters 2, 4, 6, and 7, which all have peak expression in the light phase when most of the chloroplast or photosynthesis related genes are also expressed (**Figure 4C, Figure S2**).

**Figure 4.**
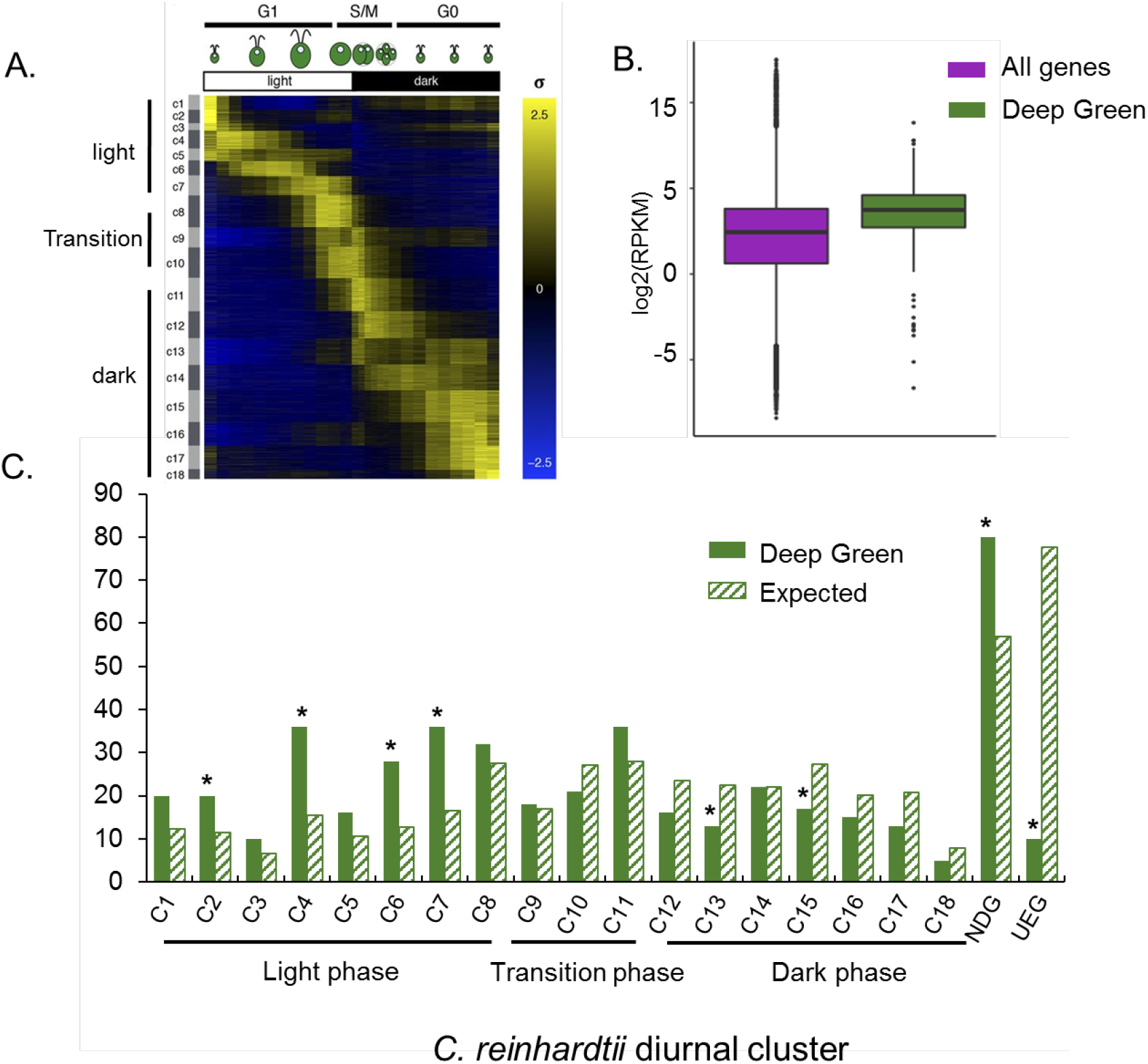
Relative expression levels and diurnal expression patterning of Deep Green genes. (A) Expression heatmap of differentially expressed genes in diurnally synchronized cultures as described previously (Zones et al., 2015, reproduced with permission from the authors). (B) Average transcript abundance of Deep Green genes in the diurnal transcriptome compared to all genes (p-value = 3.0 e-37). (C) Enrichment of Deep Green genes in 18 diurnal expression clusters shown in panel A with peak expression times shown in the same order. Non-differentially expressed (NDG), and unexpressed (UEG) groups in the diurnal transcriptome are on the right side. Significant enrichment or de-enrichment determined by Fisher’s Exact Test (background size 17,737) is indicated (*FDR corrected p-value < 0.05).

Deep Green genes were also significantly over-represented in the non-differentially expressed cluster (i.e., constitutively expressed genes) and are de-enriched in the un-expressed group. Finally, Deep Green genes were also significantly de-enriched in the dark phase clusters 13 and 15, cell motility and protein post-translation modification respectively. These results combined with the enrichment for Deep Green proteins targeted to the chloroplast (**Figure 3**), suggest that approximately 60% of Deep Green genes may have important fundamental roles in chloroplast function or biogenesis. We also examined expression of Chlamydomonas Deep Green genes in a set of data that identified genes upregulated during and recovery from heat stress (Zhang et al., 2022). Deep Green genes were identified in transcriptome data of wild type Chlamydomonas cells in response to 24 h high temperature treatments of 35°C or 40°C followed by recovery at 25°C (**Figure 5**). Of the Deep Green genes present in the RNA-seq dataset, 130 (29%) and 284 (62%) were significantly up-regulated during heat treatments (heat-induced genes, HIGs) and recovery phase (recovery-induced genes, RIGs), respectively (**Figure 5A**). Among them, 83 (18%) were significantly up-regulated during both heat treatment and recovery while only 47 (10%) were up-regulated during heat treatments but not during the recovery phase. For the Deep Green genes that were represented in the HIGs and RIGs, expression was much stronger in the 40°C treatment than in the 35°C treatment suggesting a connection between heat stress response and Deep Green gene function (**Figure 5B**). In the 40°C heat stress experiment, more than 50% of the heat inducible Deep Green genes changed their expression immediately in the first 2 or 4 h. Finally, we looked at enrichment of Deep Green genes among HIGs and RIGs from the 35°C and 40°C experiments and found significant over-representation for HIGs from both the 35°C and 40°C high temperature treatments (**Figure 5C, D**), which suggests conservation of temperature responsive genes in the green lineage. In contrast, Deep Green genes were enriched for RIGs after 40°C but not 35°C heat treatments (**Figure 5E, F**), suggesting some Deep Green genes may have potential functions in recovery from acute heat stress.

**Figure 5.**
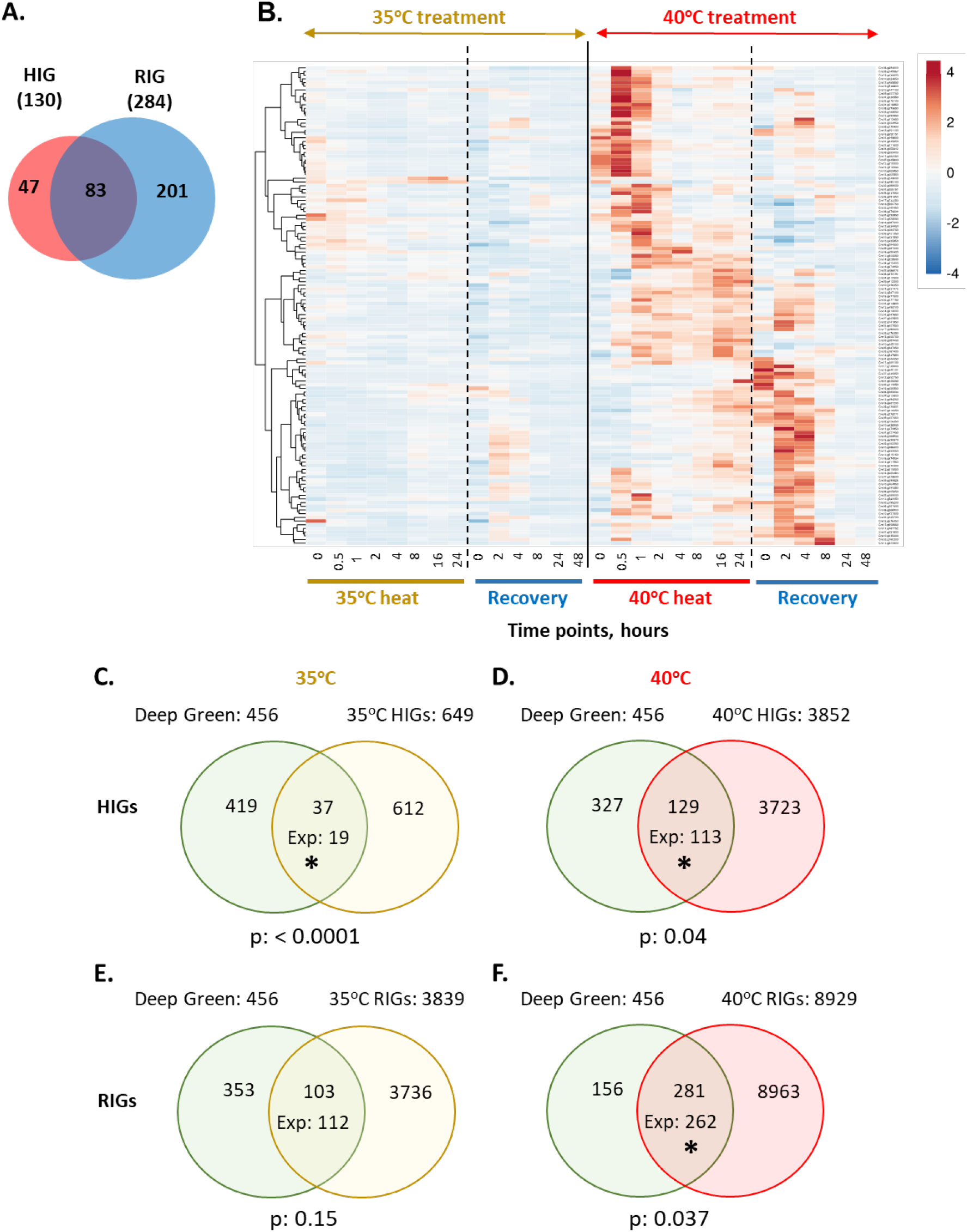
Deep Green genes are significantly enriched for heat inducible genes in Chlamydomonas. (A) Differentially expressed Deep Green genes during and after heat treatments (35°C or 40°C) described in Zhang et al., 2022. Deep Green genes with a log2(foldchange) ≥ 1 and an FDR corrected p-value < 0.05 at a minimum of one time point during heat treatment of 35°C or 40°C or during recovery were identified as heat inducible genes (HIGs) or recovery inducible genes (RIGs), respectively. (B) Heatmaps of differentially expressed Deep Green gens during and after heat treatments. Color bars represent log2(foldchange) of transcripts as compared to the preheat time point, with red colors for up-regulation and blue colors for down-regulation, white color for no differential expression. The black solid line separates the 35°C and 40°C treatments. The black dashed lines indicate the end of 24 h heat treatments. Time points indicate the length of time at the respective temperature starting from 0 hours (h) when the sample had reached the target (35°C or 40°C) or recovery (25°C) temperature. Each horizontal row represents a Deep Green gene. (C-F) Deep Green genes were significantly enriched for HIGs during (C, D) and RIGs after (E, F) heat treatments (Fisher’s Exact Test, background size 15541, *p-value < 0.05). Exp, expected overlapping numbers based on random chances. (E, F) *C. reinhardtii* Deep Green genes were significantly enriched for RIGs after 40°C heat treatment, (Fisher’s Exact Test, background size 15541, *p-value < 0.05). Exp, expected overlapping numbers based on random chances.

As a final and more general test of co-expression based functional annotation, we examined the distributions of Deep Green genes in ChlamyNET, a web-based tool to explore gene co-expression networks based on published transcriptome data (Romero-Campero et al., 2016). ChlamyNET has 9 major co-expression clusters among 9171 Chlamydomonas genes and captures co-expression relationships established under 25 different growth conditions. Among all 464 Chlamydomonas Deep Green genes, 240 are also represented in ChlamyNet. Deep Green genes were significantly enriched in clusters containing proteins associated with protein assembly and degradation and translation and lipid metabolism in the co-expression network (**Figure 6**). Both of these ChlamyNet clusters with over-representation of Deep Green genes were themselves over-represented for light phase or light-dark transition phase genes as previously defined (Zones et al., 2015, Matt et al., 2018) (**Figure S3**). Taken together, our results suggest critical conserved but unexplored functions for many Chlamydomonas Deep Green proteins in photosynthetic biology and stress responses.

**Figure 6.**
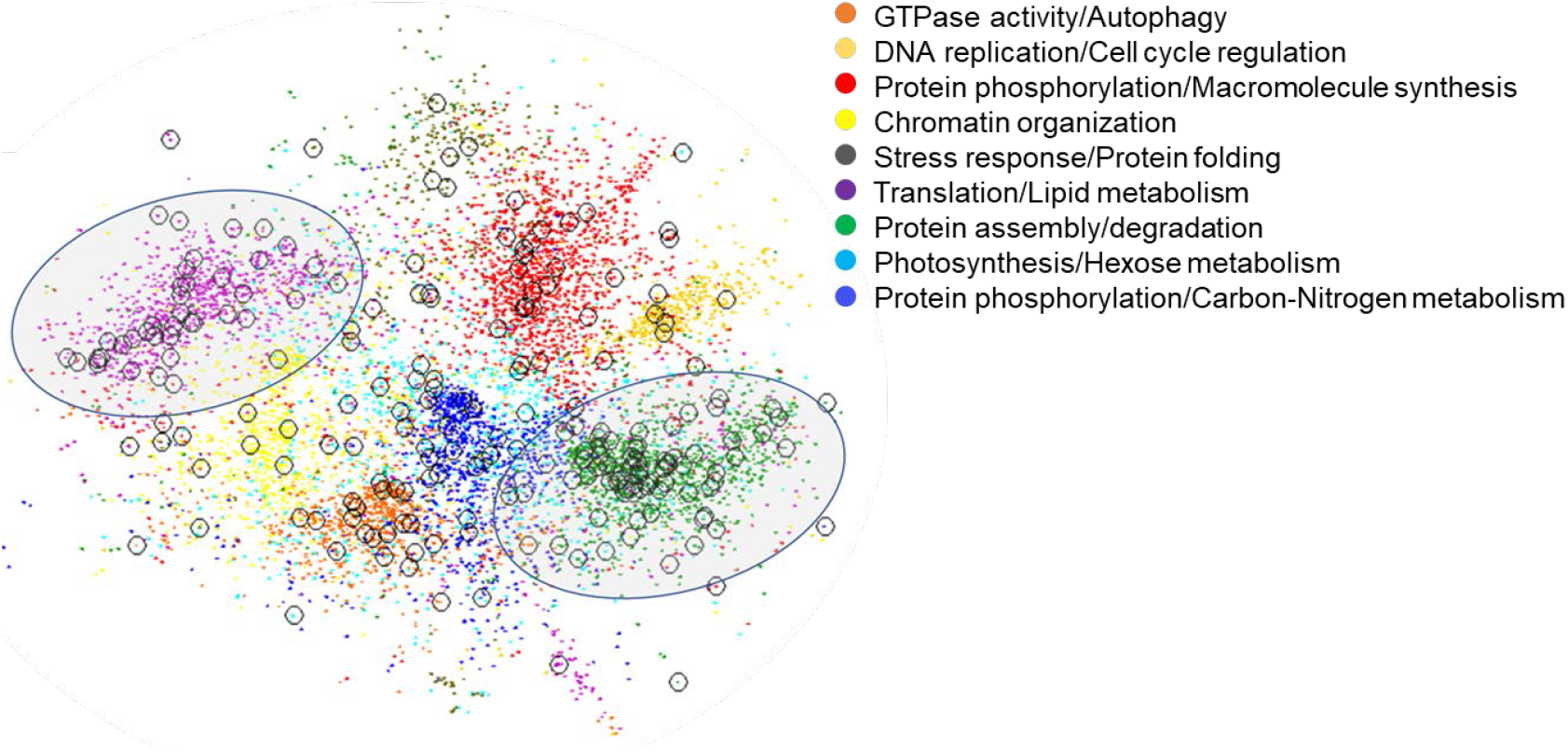
Distribution of Deep Green genes in the ChlamyNet cluster network (Romero-Campero et al., 2016). Two-dimensional graph showing gene expression clusters that are color coded, with each point representing a gene in ChlamyNet whose subcluster enrichment profile is shown in the color key. Deep Green genes are circled, and the two large subclusters with enriched representation of Deep Green genes are demarcated by shaded ovals.

### Structural properties of Deep Green proteins

Tertiary structures for each Deep Green protein were predicted using AlphaFold v2.1 and were uploaded as the Deep Green protein set to https://alphafold.ebi.ac.uk/ (Jumper et al., 2021) (**Figure 7, Table S8**). Deep Green proteins with predicted local-distance difference test (pLDDT) confidence scores higher than 50 (1338 from among the three species) were selected to perform structural matching in the Protein Data Bank (PDB) using Foldseek (van Kempen et al., 2022) (**Table S8**). Foldseek creates a template modeling (TM) score between 0 and 1 that reflects structural differences between an input query protein and its best structural match in PDB. TM scores below 0.5 indicate significantly different structures while those close to 1 are near perfect matches. The TM score distribution between the predicted tertiary structures and extant protein models in the PDB showed 777 out of 1338 (58.1%) of Deep Green proteins from all three species (268 out of 455 for Arabidopsis, 220 out of 381 for Setaria, and 289 out of 502 for Chlamydomonas) to have TM scores less than 0.5, a common threshold for protein structural comparison, and therefore are likely to have structures with novel folds (Xu et al., 2010) (**Figure 8A**). In addition, 9 of the Deep Green proteins with a potentially novel fold showed a high structural similarity among the three interspecific homologs as exemplified in (**Table 1, Figure 8B**). In addition, approximately 60% of the Arabidopsis proteome currently has either experimentally determined structures or structures through association with related proteins in the PDB (12%), with the remaining majority (48%) having been predicted using AlphaFold (Callaway, 2022). Our predicted structures of the Deep Green and unknown protein sets will likely build on this database of predicted structures.

**Figure 7.**
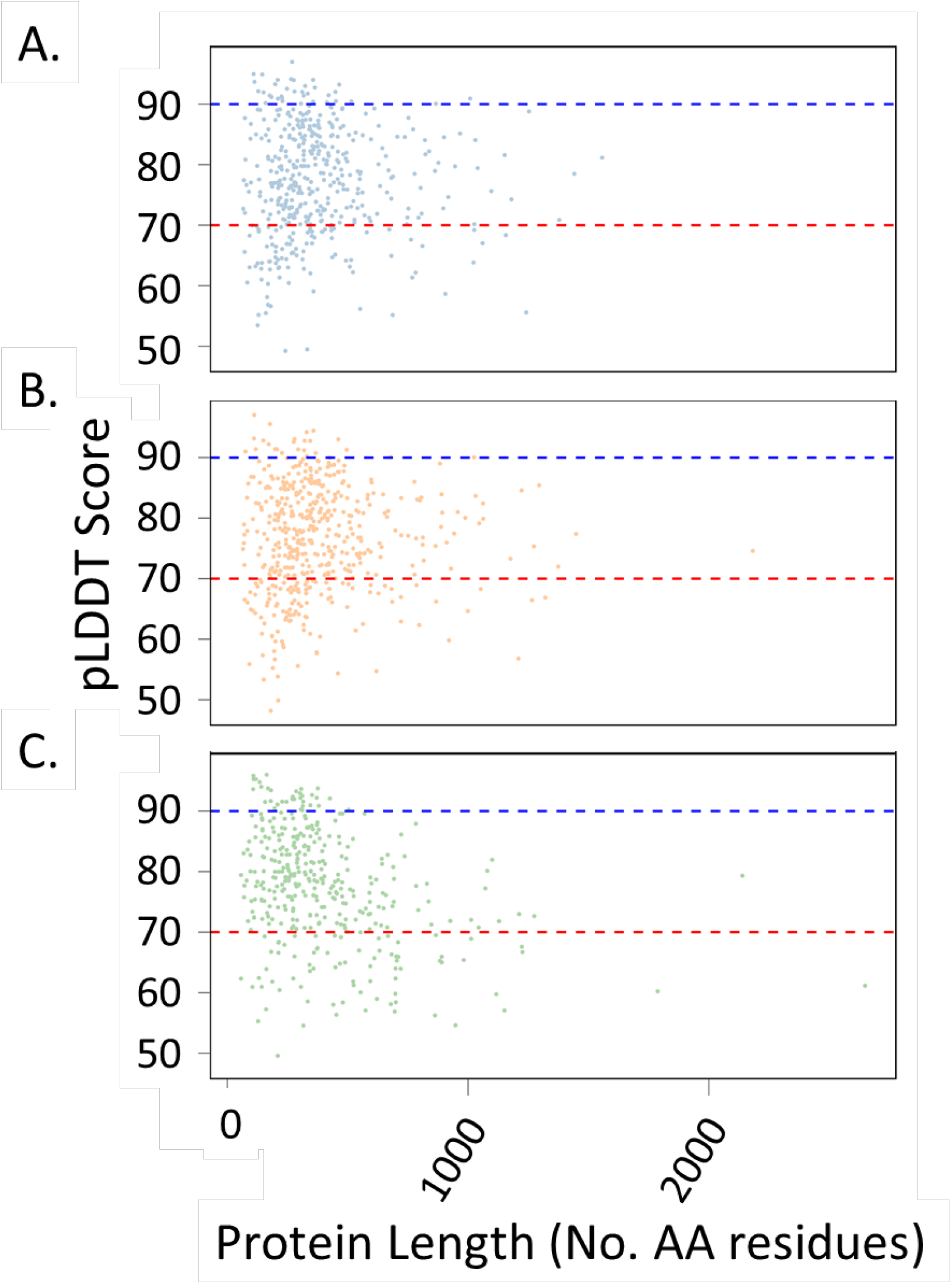
Average local-distance difference test (pLDDT) scores for Deep Green protein structures predicted using AlphaFold v2.1 for (A) Arabidopsis, (B) Setaria, and (C) Chlamydomonas. Scores >90 (blue dashed line) are considered to be highly accurate. Scores between 90 and 70 (red dashed line) are considered to indicate a generally correct backbone structure. Scores between 70 and 50 are considered to be low confidence.

**Figure 8.**
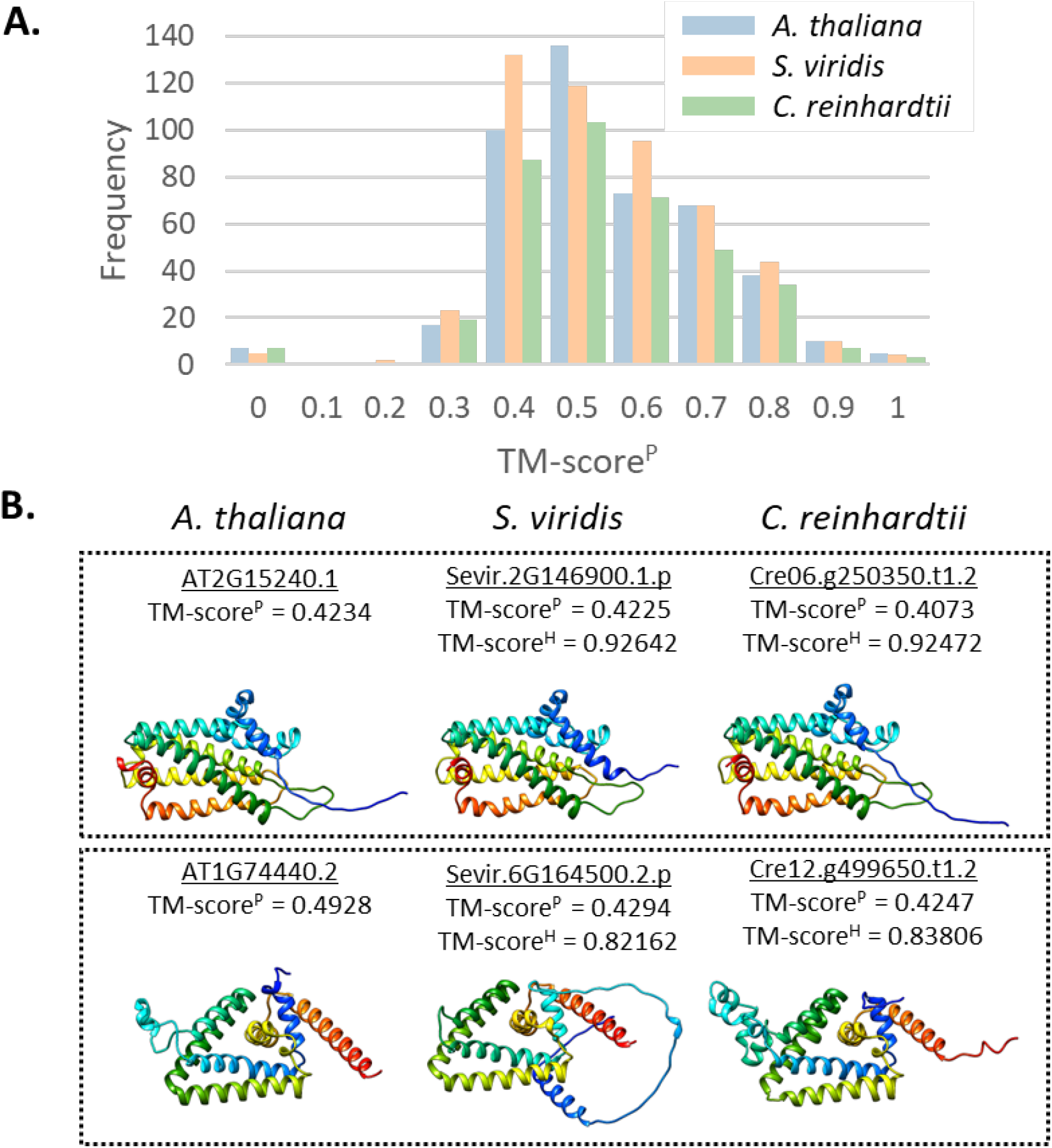
Identifying Deep Green proteins with novel structural folds. (A) TM-score^P^ distributions are shown for the best match in the Protein Data Bank (PDB) for each predicted Deep Green protein structure, with data for each species identified in the legend. A TM-score^P^ below 0.5 is considered a new fold. (B) Top and bottom boxes show structural predictions for two separate Deep Green protein families with novel structures. Ribbon diagrams show secondary structures. Best TM-scores for PDB matches are shown by TM-score^P^ and matches between Deep Green homologs are shown by TM-score^H^ compared to the Arabidopsis structure.

**Table 1.** TM-score^H^ for interspecific Deep Green protein structures having a potentially novel fold. Arabidopsis versus Setaria (Ara-Set), Arabidopsis versus Chlamydomonas (Ara-Chl), and Setaria versus Chlamydomonas (Set-Chl).

| Arabidopsis | Setaria | Chlamydomonas | TM-score<sup>H</sup> |  |  |
| --- | --- | --- | --- | --- | --- |
|  |  |  | Ara-Set | Ara-Chl | Set-Chl |
| AT2G41770.1 | Sevir.3G262100.1.p | Cre06.g260650.t1.1 | 0.708 | 0.446 | 0.487 |
| AT1G28140.1 | Sevir.9G010200.1.p | Cre04.g228450.t1.2 | 0.847 | 0.781 | 0.794 |
| AT1G74440.2 | Sevir.6G164500.2.p | Cre12.g499650.t1.2 | 0.822 | 0.838 | 0.680 |
| AT3G28720.2 | Sevir.4G242300.1.p | Cre02.g081650.t1.2 | 0.826 | 0.774 | 0.649 |
| AT5G11960.1 | Sevir.5G293000.1.p | Cre10.g419700.t1.1 | 0.913 | 0.732 | 0.650 |
| AT1G21370.1 | Sevir.7G112400.1.p | Cre02.g116000.t1.1 | 0.795 | 0.533 | 0.452 |
| AT1G04190.1 | Sevir.3G398100.1.p | Cre02.g106850.t1.2 | 0.797 | 0.532 | 0.951 |
| AT2G15240.1 | Sevir.2G146900.1.p | Cre06.g250350.t1.2 | 0.926 | 0.925 | 0.659 |
| AT3G60810.1 | Sevir.9G003500.1.p | Cre11.g468750.t1.2 | 0.722 | 0.734 | 0.681 |

To better understand the structural complexity of the Deep Green protein set, two measurements describing order versus disorder were used. IUPred3 predicts the likelihood of individual residues being in a structured region (**Figure 9A)** and the percentage alanine (A), glycine (G), and proline (P) (%AGP) in each region of a protein as a strong predictor of disorder (**Figure 9B**). The distribution of the percentage of disorder values for the Deep Green proteins in each of the three organisms indicates that the Deep Green proteins have higher representation at lower percentage disorder values and lower representation at higher percentage disorder values compared to all proteins suggesting they are overall more ordered than average (**Figure 9A**). The %AGP for the Deep Green proteins also showed a lower distribution of high %AGP suggesting less disorder among the Deep Green protein set (**Figure 9B**). We also note, as previously observed (Basile et al., 2017), a correlation between GC content in each of the three genomes and the overall amount of proteome structural disorder predicted from %AGP since the three AGP codons are represented by GC rich triplets. Chlamydomonas has the greatest amount of disorder in its predicted proteome and the highest genomic GC content (66%), while Arabidopsis has the lowest predicted disorder and the lowest GC content (36%) with Setaria in between (46% GC). Nonetheless, within each species the Deep Green predicted proteins were predicted to be more structured than average proteins.

**Figure 9.**
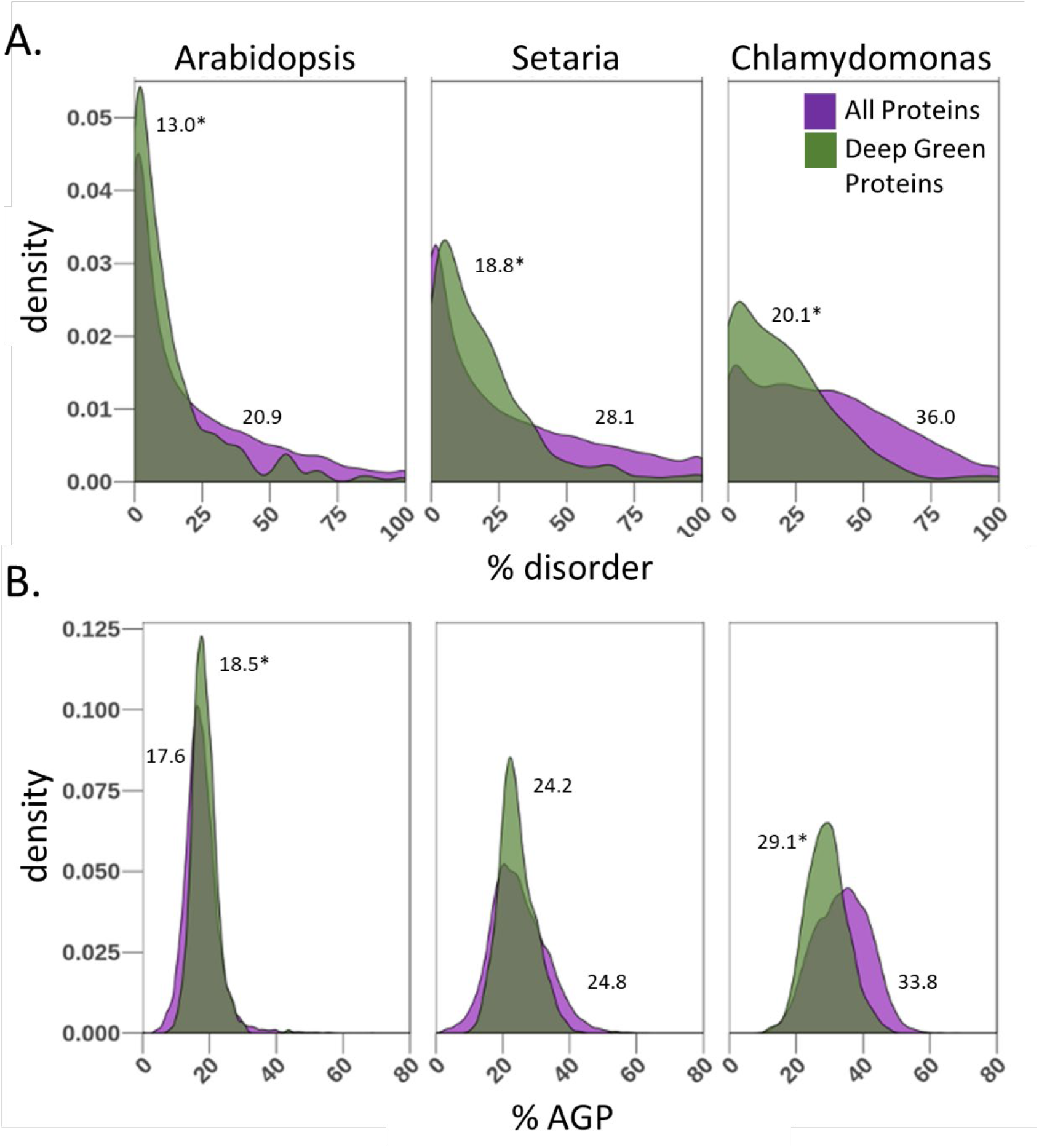
Distributions of (A) predicted % disorder or (B) % residues correlated with disorder, alanine, glycine, and proline (AGP). Each panel shows distribution of all predicted primary proteins in the indicated species and the subset of Deep Green proteins including the mean % value positioned near the respective dataset. *p-value < 0.05.

## Summary and Perspectives

The motivation behind this study was that among unknown proteins, those which are conserved in diverse members of the green lineage are likely to play important roles in photosynthetic biology. Our identification and preliminary characterization of Deep Green proteins presented here supported this hypothesis. There was a strong over-representation of Deep Green proteins for predicted chloroplast localization, suggesting their direct participation in plastid biogenesis or photosynthetic functions. In agreement with this finding, Deep Green genes in Chlamydomonas were shown to be over-represented for having light phase expression patterns that are also a characteristic of photosynthesis related proteins. Interestingly, there was also higher than expected representation of Deep Green genes among those that are induced during heat stress or heat stress recovery in Chlamydomonas, possibly reflecting the important role of chloroplast function in heat stress tolerance (Hu et al., 2020, Luo et al., 2021). The concentration of Deep Green genes in several ChlamyNet sub-networks was harder to interpret as the sub-networks contain hundreds or thousands of genes, but it again suggests some functional coherence for the Deep Green gene list whose members were not distributed uniformly within ChlamyNet.

A second goal of identifying Deep Green proteins was to investigate possible new protein folds and structures as these could also have novel catalytic or other properties. Using the powerful new version of Alphafold v2.1, we predicted stable structures for a large fraction of Deep Green proteins; and excitingly, many of them had no good matches in PDB meaning that they likely represent new families of structural folds. Also encouraging was the agreement found between predicted structures of Deep Green homologs between the three species supporting the hypothesis that there will also be structure-based functional conservation across the green lineage.

Comprehensive information on conserved plant gene function would be valuable not only for basic science to promote understanding of photosynthetic biology, but could also help in understanding the contributions of unknown genes to important agronomic traits. For example, four Setaria loci (Sevir.1G224300, Sevir.5G282600, Sevir.5G335650, Sevir.9G583700) identified as being linked to responses to extremes in precipitation and temperature (Mamidi et al., 2020), and 107 genes up- or down-regulated in response to aphid infection (Dangol et al., 2022) were part of our Setaria Deep Green protein set and are excellent candidates for further characterization.

In summary, the Deep Green gene/protein list that has been created and characterized here will be an impactful starting point for applying functional genomics and structural studies that will help shed light on unexplored areas of biology in photosynthetic eukaryotes.

## Materials and Methods

### Datasets

Current protein lists for Arabidopsis, Setaria, and Chlamydomonas were downloaded from Phytozome 13 (https://phytozome.jgi.doe.gov/pz/portal.html). The files downloaded and used in our analysis were: Arabidopsis v447_Araport11: Athaliana_447_Araport11.annotation_info.txt, Athaliana_447_Araport11.define.txt, Athaliana_447_Araport11.protein_primaryTranscriptOnly.fa; Setaria v2.1: Sviridis_500_v2.1.annotation_info.txt, Sviridis_500_v2.1.defline.txt, Sviridis_500_v2.1.protein_primaryTranscriptOnly.fa; and Chlamydomonas v5.6: Creinhardtii_281_v5.6.annotation_info.txt, Creinhardtii_281_v5.6.defline.txt, Creinhardtii_281_v5.6.description.txt, Creinhardtii_281_v5.6.protein_primaryTranscriptOnly.fa. For the Arabidopsis protein set, Araport11 was chosen over TAIR10 because it is a comprehensive re-annotation of the Col-0 genome using 113 public RNA-seq data sets and other annotation contributions from the National Center for Biotechnology Information (NCBI), Uniprot, and labs conducting Arabidopsis research (https://www.araport.org/data/araport11). The initial, primary transcript-only protein lists contained 27,654, 38,334, and 17,741 proteins for Arabidopsis, Setaria, and Chlamydomonas respectively.

### Homologs

The proteins in each organism were searched against each other organism using BLASTP to identify those proteins having homologs using both BLOSUM 45 and 62 matrices with an e-value cutoff of 10^-3^ and a qcovs score ≥40% (Altschul et al., 1990).

### Clustering

To reduce the overall number of proteins and to generate a non-redundant protein set, the three-step hierarchical clustering algorithm h3-cd-hit (http://weizhong-lab.ucsd.edu/webMGA/server/) was used on the protein list derived from the primary transcript only lists from each of the three organisms to identify those proteins that cluster together with ≥30% primary sequence identity (Huang et al., 2010).

### Protein localization prediction

To characterize the functionally unknown protein sets, analyses on the primary amino acid sequences were performed. Intra- and extra-cellular localization and signal peptide cleavage site were predicted using TargetP 2.0 (Armenteros et al., 2019), WoLF PSORT (Horton et al., 2007), and PredAlgo ((Tardif et al., 2012), only for *C. reinhardtii*), transmembrane domains were predicted using Phobius ((Kall et al., 2007), https://phobius.sbc.su.se/). Enrichment in cellular localization and transmembrane predictions of the unknown and Deep Green protein sets were performed using the hypergeometric or Fisher’s exact test.

### Co-expression analysis

Co-expression analyses were performed on Chlamydomonas Deep Green proteins using two different datasets as follows: (1) 18 diurnally expressed clusters and two unclustered groups (non-differentially expressed and non-expressed clusters) described in Chlamydomonas (Zones et al., 2015); (2) High-temperature and recovery inducible genes (HIGs and RIGs, respectively) identified previously (HIGs and RIGs are defined as transcripts that were induced for at least one time point during high temperatures and recovery, respectively) (Zhang et al., 2022); Deep Green genes that are present in this RNA-seq datasets were used for the enrichment analysis. Clustvis heat map clustering (Metsalu et al., 2015) was performed via correlation distance, completed clustering with tightest cluster first for rows and no clustering for columns. Gene co-expression networks in Chlamydomonas genes are derived from ChlamyNET (Romero-Campero et al., 2016). Graphical representation of the ChlamyNet cluster networks was performed using Cytoscape with an organic layout method (Smoot et al., 2011). This algorithm consists of a variant of the force-directed layout. Nodes produce repulsive forces whereas edges induce attractive forces. Nodes are then placed such that the sum of these forces are minimized. The organic layout has the effect of exposing the clustering structure of a network. In particular, this layout tends to locate tightly connected nodes with many interactions or *hub nodes* together in central areas of the network. The over-representation hypher.test analysis was performed using the R programming language with a significant level of 0.05.

### Structural Predictions

We used AlphaFold v2.1 to predict tertiary structures of the Deep Green proteins from their amino acid sequences. Five structural models were generated per protein and the models were ranked using the predicted local-distance difference test (pLDDT) scores (Jumper et al., 2021). The model with the highest pLDDT score was accepted as the most accurate structural prediction. Computations were carried out on NREL’s Eagle High-Performance Computing (HPC) cluster. Structural predictions for protein sequences with less than 1100 amino acid residues were run on graphics processing unit (GPU) nodes (with 16 GB Tesla V100 accelerators), while longer sequences were performed on graphics processing unit (CPU) nodes due to memory limitations. Twenty Arabidopsis, 26 Setaria, and 18 Chlamydomonas Deep Green protein sequences were predicted by SignalP-5.0 to contain signal peptides. For these sequences, the signal peptides were truncated *in-silico* prior to structure prediction. AlphaFold v2.1 structural prediction ran successfully on 457 out of 458 Arabidopsis, all 504 Setaria, and 382 out of 384 Chlamydomonas Deep Green proteins. AlphaFold v2.1 runtime errors occurred for the Arabidopsis protein AT1G21650.3 (1806 aa) and for two Chlamydomonas proteins, Cre04.g216050.t1.1 (3691 aa after removal of predicted signal peptide) and Cre07.g314900.t1.1 (732 aa). These structures could not be predicted due to runtime errors of the HHBlits software that AlphaFold v2.1 uses for fast iterative protein sequence searching by HMM-HMM alignment. AlphaFold v2.1 has been documented to fold proteins that are at least 16 and at most 2700 amino acid residues long. To perform structural homology analysis on the Deep Green proteins, proteins with a predicted tertiary structure having a confidence score higher than 0.5 were selected and FoldSeek (van Kempen et al., 2022) was used to generate structural alignments with proteins in the Protein Data Bank (PDB, version on 2021-06-01) using the parameters: --alignment-type 1 --tmscore-threshold 0 --max-seqs 2000.

### Protein disorder predictions and analyses

Protein order versus disorder based on overall secondary structure was quantified using a standalone version of the Intrinsically Unstructured Prediction (IUPred3) (Erdos et al., 2021) tool that was run on the NREL high-performance computing (HPC) cluster. In the current work, the long disorder prediction mode of IUPred3 was used along with the medium smoothing option that involves the Savitzky-Golay filter with parameters 19 and 5. IUPred3 returns a score, between 0 and 1, for each amino acid residue in the input protein sequence, that represents the probability of the given residue being part of a disordered region. Residues with scores equal to or exceeding 0.5 were considered to be disordered. Next, the percentage disorder (percentage of the total number of amino acid residues in a protein that are disordered) was quantified for each of the lead proteins from the entire proteomes of Arabidopsis, Setaria, and Chlamydomonas. The percentage disorder values for the Deep Green proteins which constitute a subset of the set of all the lead proteins were selected for additional analyses. The percentage of amino residues that are Ala, Pro, and Gly in each of the lead Arabidopsis, Setaria, and Chlamydomonas proteins were estimated as another measure of structural disorder using in-house Python code that involved use of the Biopython library (Cock et al., 2009). Signal peptides were removed using SignalP (Armenteros et al., 2019).

## Acknowledgements

This work was authored in part by the National Renewable Energy Laboratory, operated by Alliance for Sustainable Energy, LLC, for the U.S. Department of Energy (DOE) under Contract No. DE-AC36-08GO28308. Funding provided by Department of Energy’s Office of Science Biological and Environmental Research. The views expressed in the article do not necessarily represent the views of the DOE or the U.S. Government. The U.S. Government retains and the publisher, by accepting the article for publication, acknowledges that the U.S. Government retains a nonexclusive, paid-up, irrevocable, worldwide license to publish or reproduce the published form of this work, or allow others to do so, for U.S. Government purposes. This research was also partially supported by DOE award No. 0020400 to Ru Zhang. Erin Mattoon was supported by the William H. Danforth Fellowship in Plant Sciences, DDPSC start-up funding (to Ru Zhang) and Washington University in St. Louis.

## Author Contributions

EPK designed and performed the research, analyzed data, and wrote the paper. AN, PS, HN, CC, EMM, NNZ, and RZ performed research and analyzed data. PS, HN, and NNZ manually checked the Deep Green gene list and its overlap with GreenCut2. HN, and EMM investigated Chlamydomonas Deep Green genes in algal transcriptomes with high temperature treatments. JL and JC designed and implemented protein structure and fold analysis. JU designed the research and analyzed data. PSJ developed the workflow for running AlpahFold2 predictions on NREL’s HPC system. All co-authors helped revise the paper.

## Supplemental Figures

**Figure S1.**
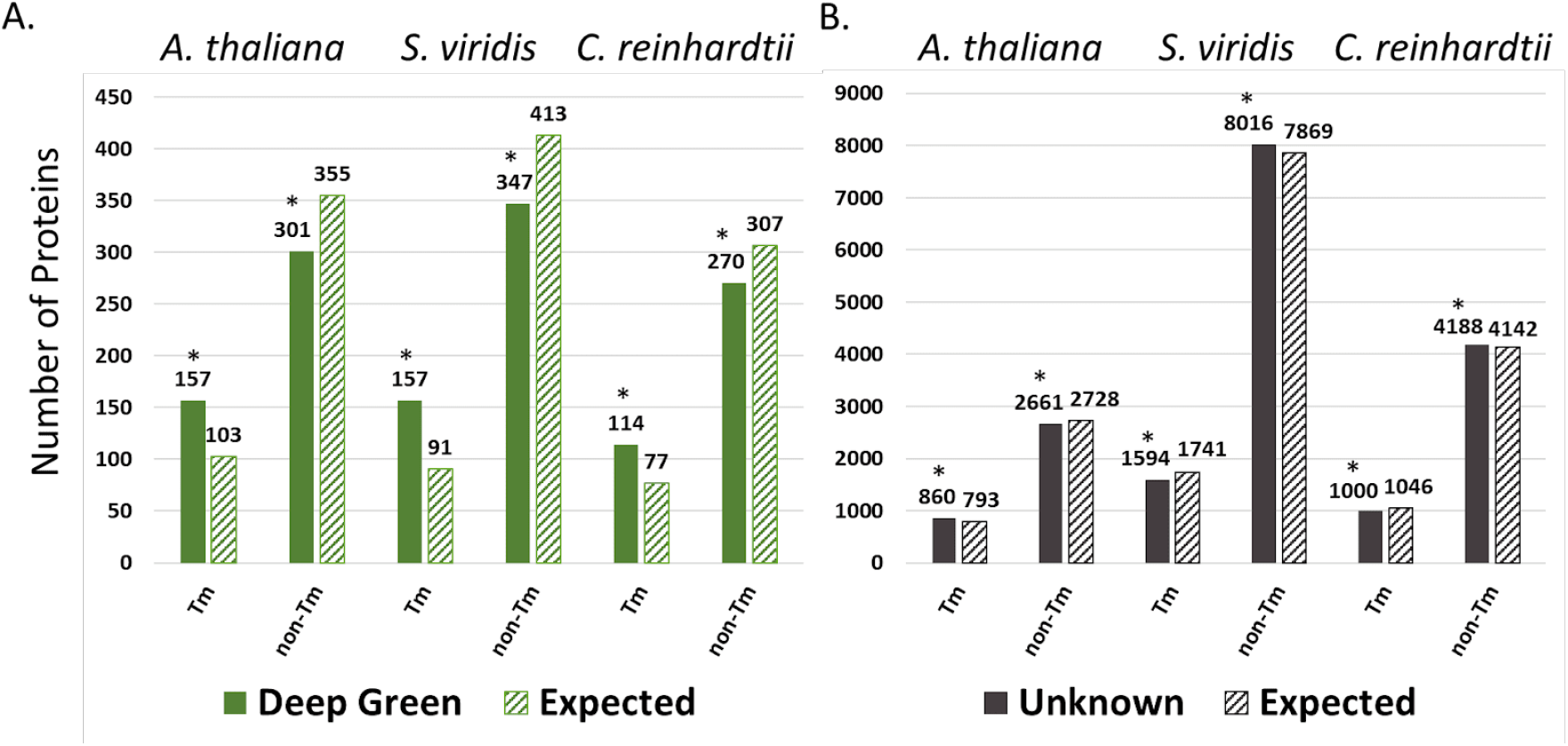
Number of proteins predicted to contain transmembrane (Tm) domains in (A) Deep Green and (B) Unknown proteins using Fisher’s Exact Test (Background size 13058, 19952, and 14076 for Arabidopsis, Setaria, and Chlamydomonas respectively) based on the total number of proteins. Significant enrichment or de-enrichment (*, FDR corrected p-value < 0.05) is indicated.

**Figure S2.**
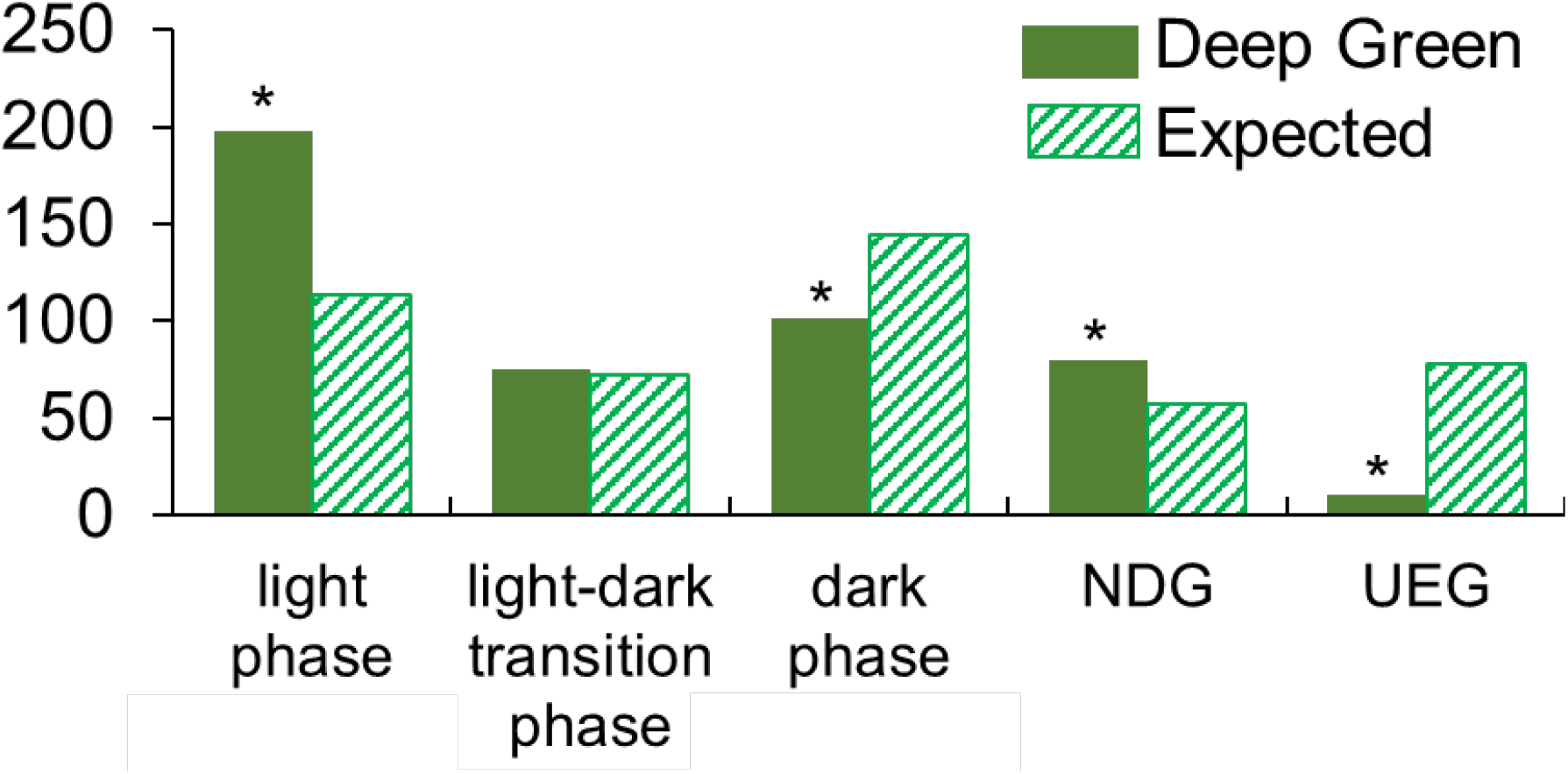
Enrichment of Deep Green genes in light phase (clusters 1-8), light-dark transition (clusters 9-11), dark phase (clusters 12-18), and non-differentially expressed (NDG) and unexpressed (UEG) groups in the diurnal transcriptome. Asterisks indicates significant enrichment or de-enrichment (* FDR < 0.05).

**Figure S3.**
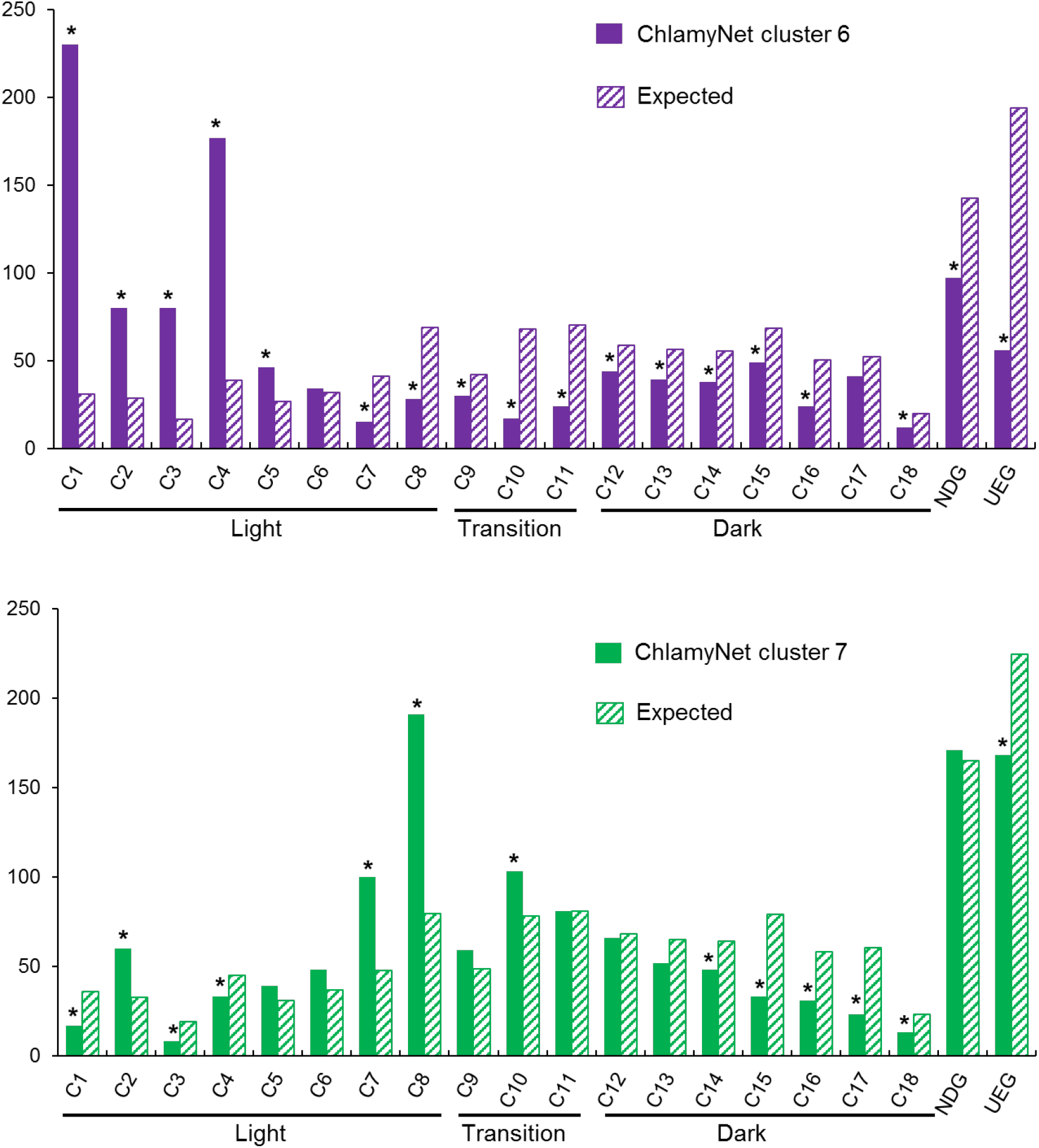
Enrichment of Deep Green genes in ChlamyNet clusters 6 and 7 in the diurnal transcriptome. (A) Enrichment of ChlamyNet cluster 6 (purple cluster) in diurnal transcriptome. (B) Enrichment of ChlamyNet cluster 7 (green cluster) in diurnal transcriptome. Fisher’s Exact Test (background size 17737) was used to determine if ChlamyNet cluster 6 and cluster 7 are enriched in diurnal transcriptome (* FDR < 0.05). NDG, non-differentially expressed group; UEG, unexpressed groups.

